# Instance segmentation of mitochondria in electron microscopy images with a generalist deep learning model

**DOI:** 10.1101/2022.03.17.484806

**Authors:** Ryan Conrad, Kedar Narayan

## Abstract

Mitochondria are extremely pleomorphic organelles. Automatically annotating each one accurately and precisely in any 2D or volume electron microscopy (EM) image is an unsolved computational challenge. Current deep learning-based approaches train models on images that provide limited cellular contexts, precluding generality. To address this, we amassed a highly heterogeneous ∼1.5 x 10^6^ image 2D unlabeled cellular EM dataset, and segmented ∼135,000 mitochondrial instances therein. MitoNet, a model trained on these resources, performs well on challenging benchmarks and on previously unseen volume EM datasets containing tens of thousands of mitochondria. We release a new Python package and napari plugin, empanada, to rapidly run inference, visualize, and proofread instance segmentations.

## Introduction

Electron microscopy (EM) reveals high-resolution snapshots of cellular and subcellular ultrastructure. Recent volume EM advances have enabled 3D imaging ^1, 2^ of increasingly large specimens, notably in connectomics ^3–5^ where deep learning (DL) algorithms are actively employed to generate wiring diagrams and enable quantitative analyses ^6–9^. Similar methods have also been used to investigate organellar structures in other systems ^10–14^, generating biological insights at unprecedented scales. The typical DL workflow for these applications is to densely annotate features in a 3D region of interest (ROI), train a model on these annotations, run inference on the full dataset, and then proofread the model output ^6, 7, 10, 11, 15, 16^. While visually impressive, these results belie a failure to generalize, meaning that segmentation quality drops dramatically when models are presented with images from unseen cellular milieus or different sample preparation or imaging protocols ^10, 11, 17^. Poor model generalization thus forces repeated cycles of annotate, train, infer, and proofread for every new project; this laborious workflow could be drastically simplified and accelerated by use of a generalist DL segmentation model.

As ubiquitous and morphologically complex organelles that play critical roles in cellular physiology and pathology ^18–26^, mitochondria provide both a stringent test and high payoff for such a model. Studies of key aspects of mitochondrial biology such as morphologies ^27–30^, networks ^31, 32^ and fission-fusion cycles ^33, 34^ would benefit from the high-resolution 3D imaging afforded by volume EM. But robust quantitation from these experiments requires precise and accurate labeling of each of the hundreds of “instances” per cell, and the solution must be efficient and general to process large and varied volume EM datasets. Fortunately, despite their extraordinary variety, mitochondria are instantly recognizable by their ultrastructure, hinting at potential for a “universal” DL model that can accurately and precisely recognize them in any EM image. However, their heterogeneity and the vast cellular landscape within which they are presented means that: 1. This task is fundamentally different from neuronal tracing, and 2. The model may not be well trained by homogenous datasets such as MitoEM ^15^, currently the largest available labeled dataset, derived from brain tissue only. We hypothesized that sparse 2D instance segmentations from an eclectic set of EM images could effectively expand the range of contexts needed for model generalization, and at a much lower cost than dense 3D segmentation. Here, we curated *CEM1.5M*, a heterogeneous, non-redundant, information-rich, and relevant unlabeled EM image dataset (at ∼1.5 x 10^6^ images, the largest of its kind to our knowledge) for use as a database to pre-train and sample images for mitochondrial, or other organelle, segmentation. Combining existing labeled datasets and crowdsourced annotations of images from CEM1.5M, we have created *CEM-MitoLab*, a similarly diverse dataset for training mitochondrial segmentation models. We show that *MitoNet,* the model trained on CEM1.5M and CEM-MitoLab, outperforms similar models trained on other candidate datasets when tested on new and challenging benchmarks. Finally, we use *empanada,* a new Python package and napari plugin for model training, inference, and proofreading to efficiently create tens of thousands of high-quality mitochondrial segmentations from public volume EM datasets of mouse liver and kidney tissue and make visual and quantitative comparisons between them.

## Results

### CEM1.5M and CEM-MitoLab dataset creation

Our overall curation process is outlined in Figure 1a. We first generated a database of unlabeled EM images of cells and tissues comprising 466 volume EM datasets (356 in-house, 110 externally generated and publicly deposited, total > 2 PB). To limit overrepresentation, datasets larger than 5 GB were cropped into random 3D ROIs while smaller datasets were retained as-is. This yielded 15,152 3D ROIs equaling 338 GB. To these we added 28 videos of EM stacks from online publications and 12,658 traditional 2D TEM and STEM images (5,657 in-house and NCI EM data; 7,001 external). Metadata and attribution were recorded, where possible, for all datasets (**Supplementary File 1).** We then applied our previously developed data preparation and curation pipeline ^35^ to sample xy, xz and yz planes of isotropic volumes, remove near-duplicate patches within datasets, and triage “uninformative” patches as classified by a neural network. This process created 1,592,753 unique 2D image patches, called CEM1.5M.

**Figure 1:**
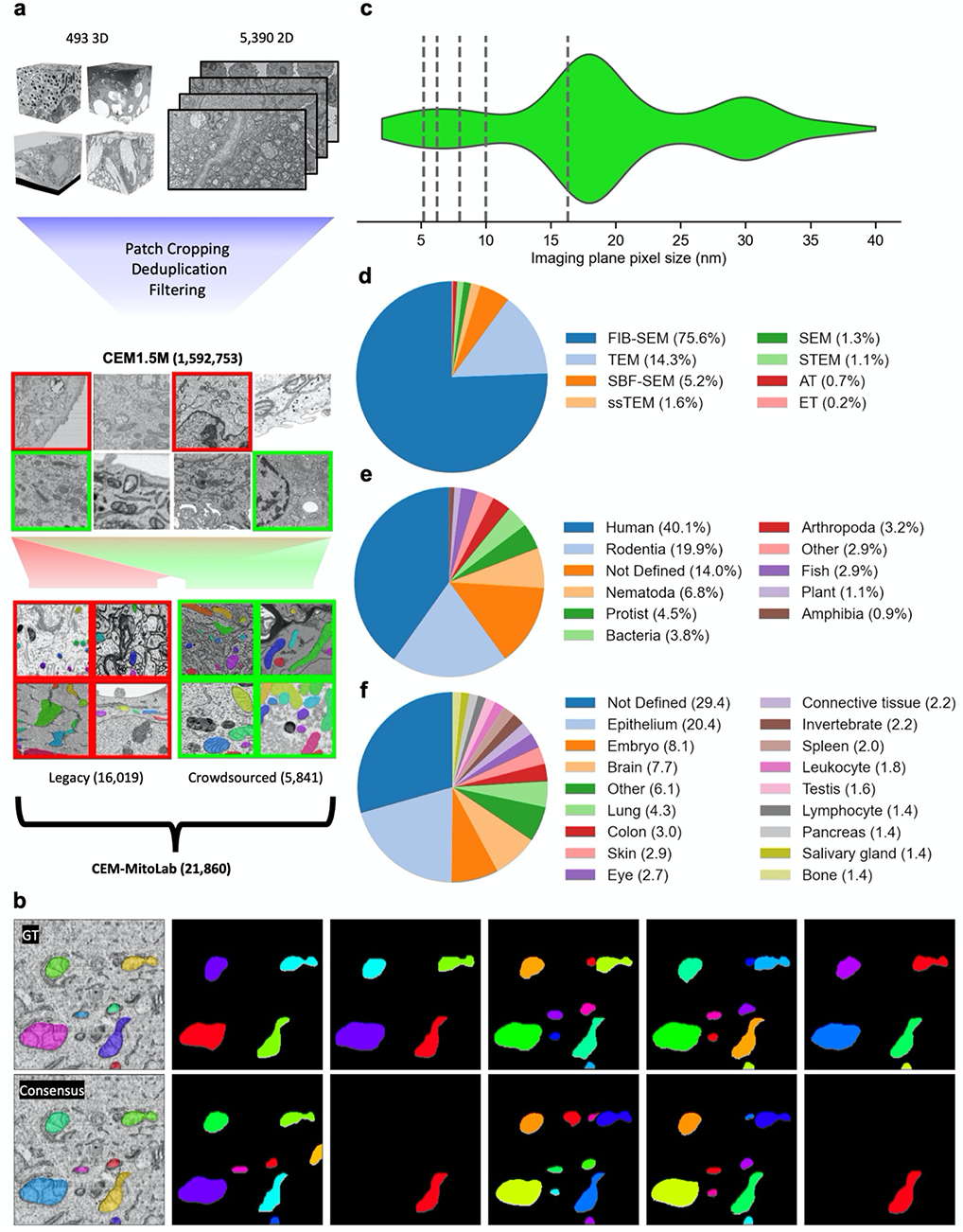
Creation of a diverse and representative dataset for mitochondrial instance segmentation. **a.** Schematic of the data curation pipeline. Volume EM reconstructions and 2D EM images were curated to create CEM1.5M. Random patches from previously labeled data (legacy annotations, red) and crowdsource annotated patches from CEM1.5M (green) were combined to form CEM-MitoLab. **b-e.** Dataset distribution by various parameters in CEM-MitoLab. **b**. Imaging plane pixel sizes of volume EM images (n=489). Dashed lines, 2D EM images. **c.** Imaging technique, **d.** Source organism, **e.** Source tissue (vertebrates only, *in vitro* cells grouped under Not Defined; n=593). **f.** Example of crowdsourced annotation with ground truth (GT, top left), consensus annotation (bottom left) and ten independent student annotations of an image showing high degree of consensus.

Of the combined 494 3D and video datasets, we selected 489 for annotation, excluding volumes with severe artifacts or > 40 nm pixel size. 12,622 cropped 2D patches from 19 of the volumes with mitochondrial instances already segmented (7 in-house, 12 external) were combined with 3,397 other 2D in-house segmentations to form the “legacy” dataset. Inspired by the citizen science project Etch-a-cell ^13^, we used the Zooniverse platform (Supplementary Figure 1a) to crowdsource the annotation of 5,481 additional images to 34 briefly trained local high school students. To evenly represent the collected data, we sampled a maximum of 15 random patches from every 3D dataset and up to 25 patches from each of 83 directories containing 2D datasets (Supplementary Figure 2a). Each image was annotated independently by ten students (Figure 1b, Supplementary Figure 1b) and we developed a robust algorithm to combine these annotations into a consensus instance segmentation (Supplementary Figure 3). All consensus segmentations were reviewed by at least two experienced researchers to create the final ground truth. The total 21,860 annotated images constitute *CEM-MitoLab* and contain 135,285 mitochondrial instances. They represent myriad image sizes, pixel resolutions, imaging techniques, sample preparation protocols, cells, tissues, and organisms (Figure 1c-f, Supplementary Figure 2b, c). Our dataset consists exclusively of 2D images and favors a broad but superficial sampling of numerous cellular contexts in comparison to the more homogenous appearance of mitochondria in singly-sourced datasets ^10, 15^ (Supplementary Figure 2d, e).

### Benchmark dataset characterization

To assess how well models trained on CEM1.5M and CEM-MitoLab generalize to unseen images, we withheld six volumes from the above pipelines and created mitochondrial segmentations for each. These included conventionally fixed, heavy metal stained, and resin-embedded samples of fly brain ^36^, mouse brain ^37, 38^, glycolytic muscle ^32^, mouse salivary gland, and HeLa cell in addition to a high-pressure frozen (HPF), freeze substituted C. elegans, all independently imaged by FIB-SEM and with isotropic voxel sampling. The first three were externally generated and the last three in-house. The mouse brain dataset was the well-studied Lucchi++ benchmark test set with the original semantic segmentations converted to instance segmentations. (Note, the Lucchi++ training set was excluded from CEM1.5M and CEM-MitoLab). Lucchi++ has 5 nm voxels and we intentionally chose or resampled the other volumes to 10-25 nm resolutions (fly brain 12 nm, HeLa 15 nm, salivary gland 15 nm, glycolytic muscle 18 nm, C. elegans 24 nm); sufficient to easily identify mitochondria and in line with the most common resolutions in the training dataset.

These benchmark volumes encapsulate different cellular contexts, mitochondrial morphologies, sample preparation protocols, and importantly, varying levels of difficulty. The simpler mitochondria in the fly brain volume have distinct variants: lightly stained with poorly defined cristae and darkly stained with well-defined cristae (Figure 2a, orange and blue arrows respectively). The HeLa cell volume is cluttered with organelles and vesicles and has localized heavy metal precipitates (green arrow). The C. elegans volume has overall lower contrast, as expected for an HPF sample, and presents difficult to segment mitochondrial morphologies – small puncta and skinny tubules (Figure 2b). There are also membranous organelles ^39^ that appear similar to mitochondria with swollen cristae at these resolutions (yellow arrow). The glycolytic muscle volume has a relatively uncluttered background but complex and elaborate branched morphologies. The salivary gland volume is the most challenging because of flat and bowl-shaped mitochondria tightly pressed against salivary granules (red arrow), weak staining, and close packing of the mitochondria themselves (Figure 2b).

**Figure 2:**
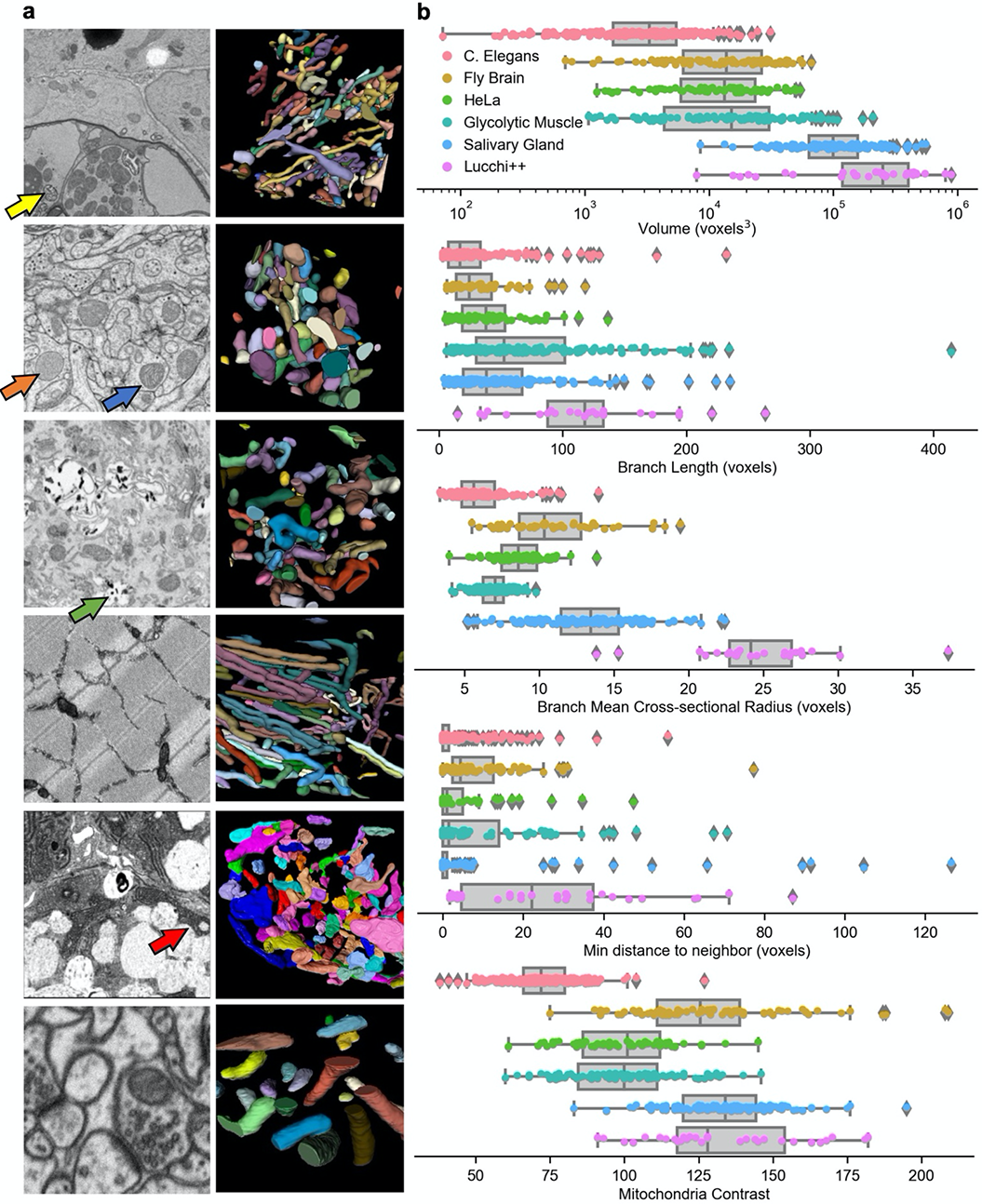
Challenging and diverse benchmarks for evaluating automatic instance segmentation performance. **a.** 2D representative images (left) and 3D reconstructions (right) for the benchmark test sets. Top to bottom: C. elegans, Fly brain, HeLa cell, Glycolytic muscle, Salivary gland, Lucchi++. Yellow arrow, membranous organelle; orange and blue arrows, lightly and darkly stained mitochondria; green arrow, heavy metal precipitate; red arrow, mitochondrion and tightly apposed salivary granule in the acinus. **b.** Comparison of individual mitochondria and box plots across benchmarks by (top to bottom): volume (log scale), branch length, mean cross-section radius, minimum distance to neighbor (all in voxels) and mitochondrial contrast. Pink, C. elegans n=241; yellow, Fly brain n=91; green, HeLa cell n=68; teal, Glycolytic muscle n=104; blue, Salivary gland n=131; purple, Lucchi++ n=33.

### Deep learning model and postprocessing

For 2D instance segmentation, we adopted Panoptic-DeepLab ^40^ (PDL). PDL is an encoder-decoder architecture, like the standard U-Net ^41^, that takes a bottom-up approach to instance segmentation and scales to segment an arbitrarily large numbers of objects within a field of view (top-down and direct set prediction algorithms like Mask R-CNN ^42^ and Mask2Former ^43^ cap the number of detections). We trained PDL models to infer mitochondrial semantic segmentations, centers and per-pixel x and y offsets from the associated object’s center (Figure 3a). To create segmentations at resolutions higher than the input image resolution, we employed PointRend ^44^, which iteratively interpolates and reevaluates the label of the most uncertain pixels in a segmentation (Supplementary Figure 4). This feature allows the model to accept downsampled images and output precise segmentation boundaries at the original resolution without a resolution-specific model architecture. The semantic segmentation, offsets, and object centers were postprocessed into an instance segmentation.

**Figure 3:**
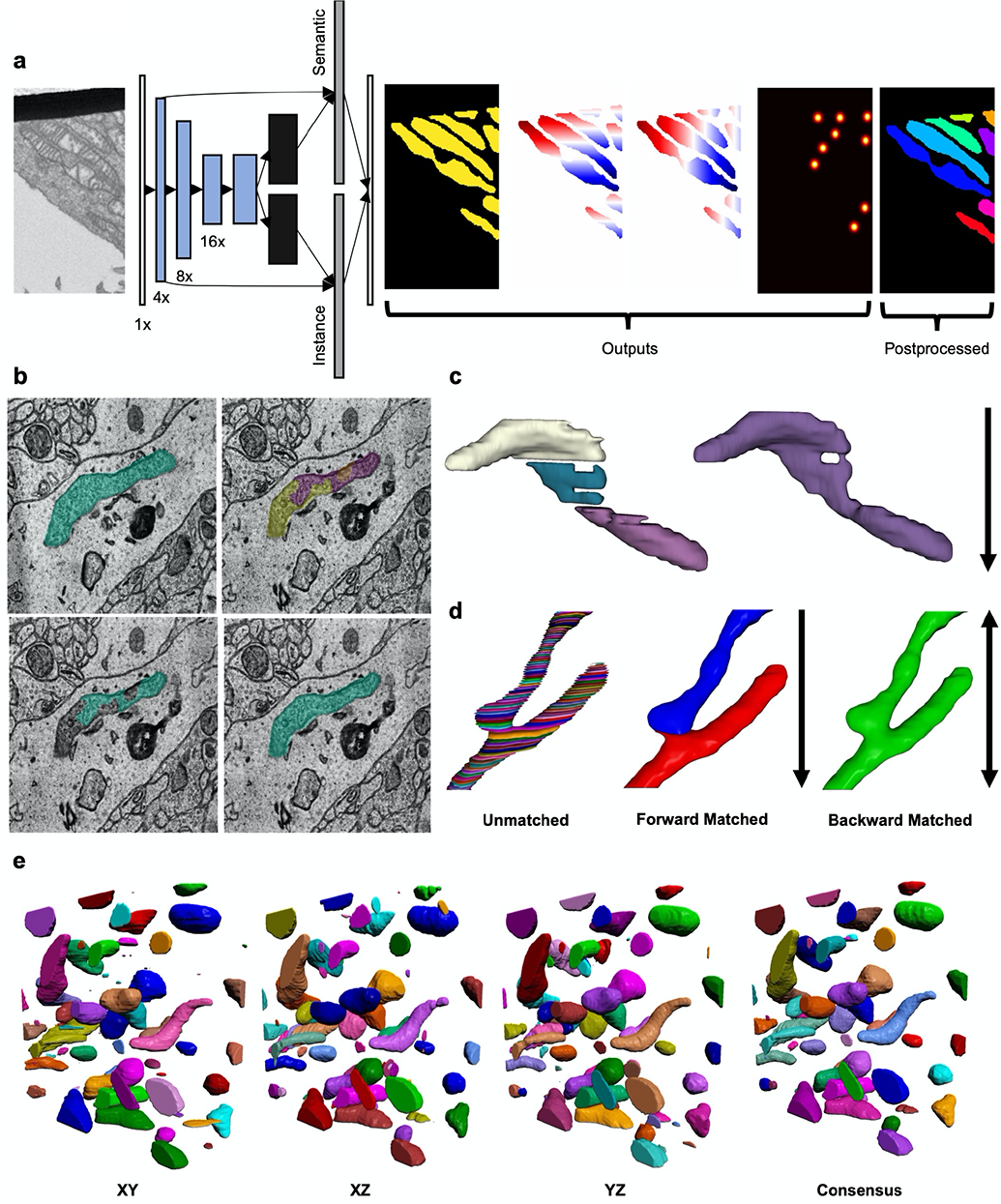
Deep learning model and postprocessing pipeline to create 2D or 3D instance segmentations. **a.** Schematic of Panoptic-DeepLab showing the input grayscale image (left;, blue boxes, encoder layer outputs; black boxes, ASPP layer outputs; gray boxes, decoder layer output. Outputs of the network are (left to right) semantic segmentation, up-down offsets, left-right offsets, and the instance centers heatmap. Far right, instance segmentation created from the outputs. **b.** Instance matching across adjacent slices uses intersection-over-union (IoU) and intersection-over-area (IoA) scores. Clockwise from top left: predicted segmentation of slice j, j+1, IoU and IoA merging, IoU only merging. **c.** Result of median filtering in direction of black arrow **d.** From left to right: Stacked 2D segmentations of before matching, after forward matching only and after forward and backward matching. Black arrows denote direction of matching. **e.** An example of 3D instance segmentation of mitochondria after running inference in (left to right) xy, xz, yz directions, and far right, and merging them into a consensus.

To track 2D instance predictions through a volume EM stack, we performed 1-to-1 matching of objects across consecutive slices using the Hungarian algorithm ^45^. We also adopted two postprocessing steps to counter oversplitting errors common for this approach. First, we computed intersection-over-area (IoA) scores for objects that were unmatched because of the 1-to-1 assumption. Unmatched objects with IoA scores over a threshold value were merged to the label of the object with which they overlapped in the preceding slice (Figure 3b). Second, we calculated 2D instance segmentations using median semantic segmentation probabilities over neighborhoods of three or more slices to cover gaps and produce smooth and continuous objects in 3D (Figure 3c). We then performed this matching in the reverse direction through the stack to account for branched morphologies (Figure 3d). Optionally, for isotropic-voxel volumes, *ortho-plane inference* ^46^ can be applied. Here, inference is performed independently on stacks of xy, xz, and yz images and the predictions are combined into a single consensus segmentation using the same algorithm developed for our crowdsourced annotations (Figure 3e). IoA-based merging, median filtering, and ortho-plane inference all yielded significant improvements on the benchmark datasets (Supplementary Table 1).

### MitoNet training and evaluation

The encoder network in our experiments was a ResNet50 ^47^ pre-trained on CEM1.5M using the SwAV ^48^ self-supervised learning algorithm (the effect of pre-training is summarized in Supplementary Table 2). Our best performing PDL model, *MitoNet*, was trained for 120 epochs on CEM-MitoLab. On the benchmark datasets we observed that it segmented a variety of mitochondrial instances accurately in 2D and 3D (Figure 4a, b). MitoNet excelled on the relatively simple brain tissue datasets (fly brain and Lucchi++) with both semantic IoU and F1@75 scores around 0.9 or better (Figure 4c). Detailed performance metrics are included in Table 1. Strikingly, MitoNet’s F1@75 score of 0.88 on the Lucchi++ test set matched specialist models that were trained exclusively on the Lucchi++ training data ^49^.

**Figure 4.**
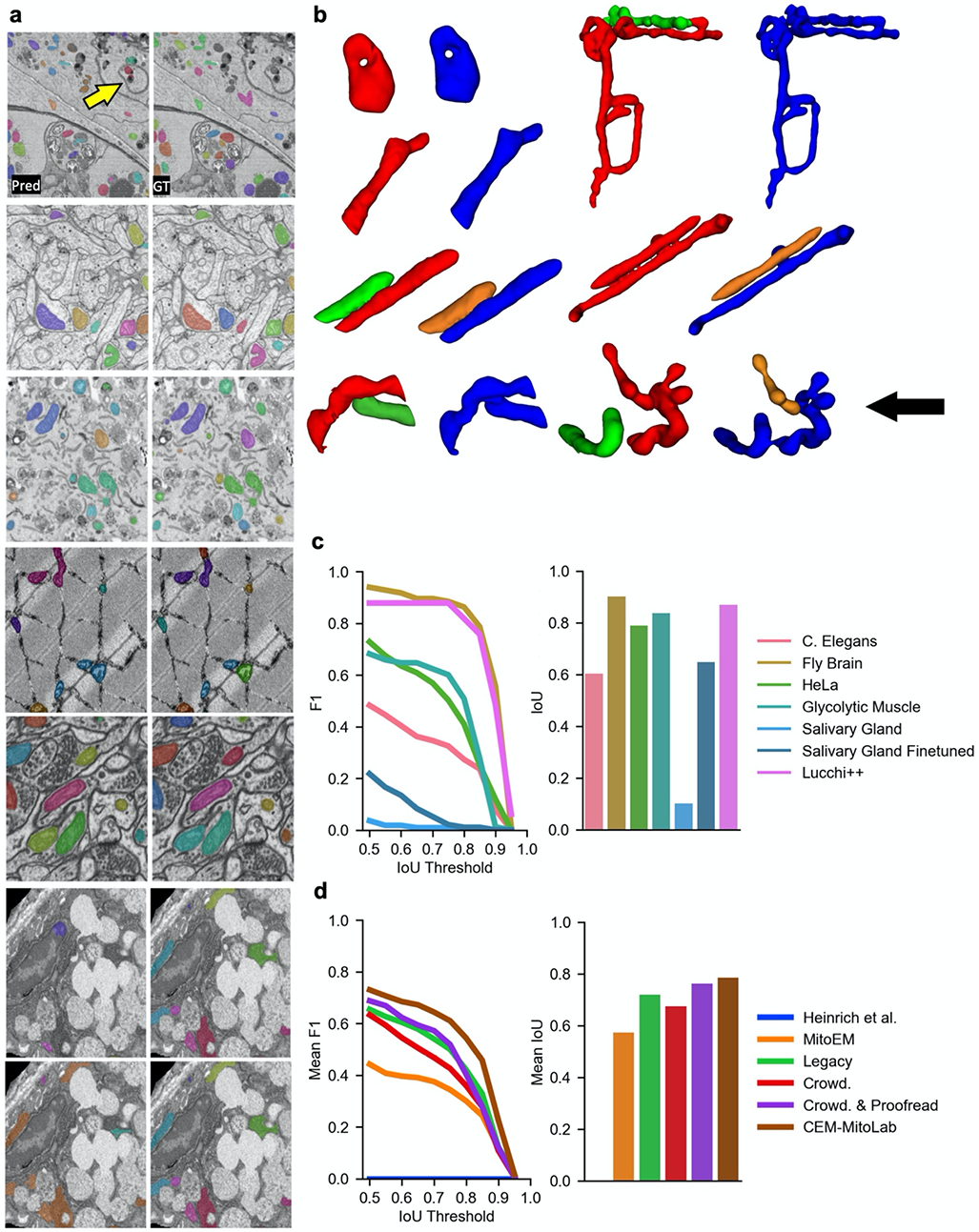
MitoNet results on benchmarks. **a.** Representative 2D images showing MitoNet segmentation performance; left column shows predictions (Pred.) and right column shows ground truth (GT). Top to bottom: C. elegans, Fly brain, HeLa cell, Glycolytic muscle, Lucchi++, Salivary gland without finetuning, Salivary gland with finetuning. **b.** Representative 3D ground truth and predicted segmentations from MitoNet. Red and green, predicted mitochondrial instances, blue and orange, ground truth instances. Black arrow, example of segmentation expected to return a high IoU but low F1 score. **c.** Left, MitoNet F1 score on each of the benchmarks as a function of IoU threshold; right, IoU scores. **d.** Left, comparison of mean F1 score for models trained on different datasets plotted against IoU threshold; right, mean IoU scores. All benchmarks except the salivary gland are included. Crowd., crowdsourced.

**Table 1:** Performance metrics of MitoNet across benchmarks. IoU, Intersection-over-Union, AP, Average Precision, PQ, Panoptic Quality.

| Dataset | IoU | F1@50 | F1@75 | AP@50 | AP@75 | PQ |
| --- | --- | --- | --- | --- | --- | --- |
| C. Elegans | 0.604 | 0.483 | 0.325 | 0.318 | 0.194 | 0.376 |
| Fly Brain | 0.902 | 0.940 | 0.885 | 0.887 | 0.794 | 0.831 |
| HeLa | 0.791 | 0.728 | 0.503 | 0.573 | 0.336 | 0.571 |
| Glycolytic Muscle | 0.840 | 0.682 | 0.601 | 0.518 | 0.430 | 0.557 |
| Salivary Gland | 0.103 | 0.036 | 0.008 | 0.018 | 0.004 | 0.022 |
| Salivary Gland Finetuned | 0.650 | 0.218 | 0.021 | 0.122 | 0.010 | 0.138 |
| Lucchi++ | 0.871 | 0.879 | 0.879 | 0.784 | 0.784 | 0.783 |

MitoNet performance on the more difficult HeLa and glycolytic muscle benchmarks was also strong with semantic IoU scores of 0.79 and 0.84, but F1@75 scores slightly lower at 0.5 and 0.6. For the HeLa benchmark, the model successfully ignored much of the cluttered intracellular features, but we did observe several FPs caused by Golgi (data not shown). The IoU and F1 scores on the glycolytic muscle dataset were surprisingly high given that only a handful of training images were from muscle tissue, and none were from glycolytic muscle (whose mitochondrial ultrastructure is significantly different ^32^). Although MitoNet correctly detected 78% of closely apposed mitochondria (i.e., < 5 voxels to nearest neighbor) at an IoU threshold of 0.5 in the HeLa volume and 52% in the glycolytic muscle, visual inspection and the lower instance compared to semantic segmentation scores suggest that identifying individual mitochondria in crowds remains a challenge (Supplementary Figure 5). Figure 4b shows examples of accurate mitochondrial segmentations, including a heavily branched mitochondrion, as well as overmerging and oversplitting errors that our method was prone to make; such errors could significantly lower F1 scores even when pixel-level accuracy was high (black arrow).

The C. elegans dataset was challenging because of small and low-contrast mitochondria that were tightly packed together. Here, we used a lower vote threshold of only one plane for ortho-plane inference to increase IoU from 0.44 to 0.60 and F1@75 from 0.18 to 0.33 (Supplementary Table 1). MitoNet struggled to detect mitochondria smaller than 1,000 voxels (∼ 0.25 µm^3^); these accounted for about 25% of all FNs (Supplementary Figure 5). The model also performed slightly worse on mitochondria with lower contrast (mean grayscale of TP, 75.9; FN, 70.5, p<0.001; see Supplementary Table 3 for complete statistics). Closely apposed instances accounted for over 75% of all FNs at an IoU threshold of 0.5. The model mostly avoided erroneously labeling the membranous organelles, but occasionally labeled some vesicles as mitochondria (Figure 4a, yellow arrow). On the most challenging benchmark, the salivary gland, MitoNet achieved an IoU score of just 0.10 and F1@50 of 0.04. Therefore, here we tested the ability of the model to adapt with minimal finetuning. We extracted 64 random patches from the ground truth data (∼0.2% of the total volume) and after training for just 500 iterations achieved dramatically improved IoU scores of 0.65 and F1@50 of 0.22 (Figure 4d). This underscores an important point: Even when the generalist MitoNet model fails on a dataset, it is still a strong starting point to train a specialist model with a modest number of examples and compute time. The low F1 score relative to IoU is because – uniquely for this dataset – a large subset of the mitochondria were tightly clustered, with touching membranes. The model and postprocessing merged nearly all of these into a single object (Figure 4a, bottom left).

Finally, we directly probed the efficacy of CEM-MitoLab against other training datasets by measuring the performance of models across all benchmarks except the salivary gland volume. We trained models for approximately 10,000 iterations to account for the different number of images in each dataset (for comparison, the 120 epochs used for MitoNet was equivalent to about 40,000 iterations). The model trained on the Heinrich et al. dataset performed poorly on the benchmarks, likely because of its small size and limited breadth. Training on MitoEM, a large but homogeneous neuronal dataset, gave F1@75 scores of 0.86 and 0.75 on the fly brain and Lucchi++ volumes, but scores of 0, 0.07, and 0.01 on the C. elegans, HeLa cell, and glycolytic muscle volumes. The model trained on expert corrected crowdsourced annotations was almost matched by a model trained on our legacy dataset – a 5x larger, reasonably heterogeneous sampling of 19 volumes, over 500 TEM images, and over 2,000 randomly chosen 2D patches from CEM500K. In both cases, IoU and F1 scores were substantially better across all benchmarks than training on MitoEM (Figure 4d, Table 2). Remarkably, even training on the *uncorrected* student consensus annotations was better than training on MitoEM for all benchmarks except the fly brain volume, emphasizing how critical data diversity is, even at the expense of per-instance accuracy (**Supplementary File 2**). Sparsely sampled, noisy, but heterogeneous labeled data can be more effective for training generalist segmentation models than much larger densely annotated, expert proofread, but homogeneous data. Unsurprisingly, our best results were achieved by training on CEM-MitoLab (i.e., the combination of legacy and expert proofread crowdsourced annotations).

**Table 2:** Average performance of MitoNet versions pre-trained on CEM1.5M but trained on different candidate datasets.

| Training Dataset | Mean IoU | Mean F1@50 | Mean F1@75 | Mean AP@50 | Mean AP@75 | Mean PQ |
| --- | --- | --- | --- | --- | --- | --- |
| Heinrich et al. | 0.000 | 0.000 | 0.000 | 0.000 | 0.000 | 0.000 |
| MitoEM | 0.574 | 0.443 | 0.344 | 0.360 | 0.286 | 0.361 |
| Legacy | 0.720 | 0.654 | 0.500 | 0.525 | 0.390 | 0.531 |
| Crowdsourced | 0.675 | 0.633 | 0.427 | 0.512 | 0.334 | 0.497 |
| Crowdsourced & Proofread | 0.763 | 0.688 | 0.519 | 0.556 | 0.388 | 0.550 |
| CEM-MitoLab | **0.786** | **0.730** | **0.610** | **0.614** | **0.493** | **0.611** |

### Analyzing Mitochondrial Morphologies in Mouse Kidney and Liver Tissue

Mitochondria are enriched in and critical to kidney function especially in proximal tubules, and their dysfunction marks renal disease and injury ^24–26, 50^. Despite the centrality of mitochondria in renal pathology, existing studies are limited by a reliance on light microscopy, 2D EM, or semantic segmentations of limited volume EM data ^51, 52^. To demonstrate the power of MitoNet for general mitochondrial instance segmentation, and to generate 3D ultrastructural correlates of known differences in mitochondrial energetics between proximal and distal tubules ^53^ we used the exact same model to automatically segment tens of thousands of mitochondria in two previously unseen volumes from mouse kidney tissue downloaded from OpenOrganelle ^36^. The ROIs contained primarily distal tubules and proximal tubules (10 GB each). Additionally, an ROI of hepatocytes from liver tissue (30 GB) were cropped from another volume for comparison. The full python package that we created to run MitoNet inference as well as the postprocessing pipeline is called empanada, while a user-friendly napari plugin of the same name allows easy GUI-based inference and post-processing, plus point-and-click merge and split operations. MitoNet segmented 7,718, 15,180, and 61,244 objects in each ROI, respectively. On a system with a single GPU and just 16 GB of RAM, automatic segmentation of all objects with empanada took 1.5 hours for each of the kidney volumes and 3.5 hours for the liver volume. Out-of-the-box, MitoNet predictions were immediately useful for visualizing the data and identifying dramatically different mitochondrial morphologies (Figure 5a). Mitochondria in both kidney volumes were densely packed into rows, polarized towards basolateral surfaces, and were revealed to have flattened morphologies. Some 6% of mitochondria in the proximal tubule (middle row) were fused into large and complex networks in sharp contrast to the much less branched morphologies in the distal tubule. Thus limited morphological assessments from light microscopy imaging of proximal tubules ^52^ can now be extended and quantified at an individual organelle level with the resolution of volume EM combined with MitoNet segmentation. Mitochondria in the hepatocytes appeared to be relatively simple tubes and spheroids; however, many instances were visible with long and skinny protrusions – possibly nanotunnels ^54^ – extending into the surrounding cytoplasm. As further evidence of the model’s ability to generalize, we observed accurate segmentations of mitochondria in endothelial cells captured by the ROIs as well (**Supplementary Video 1**).

**Figure 5.**
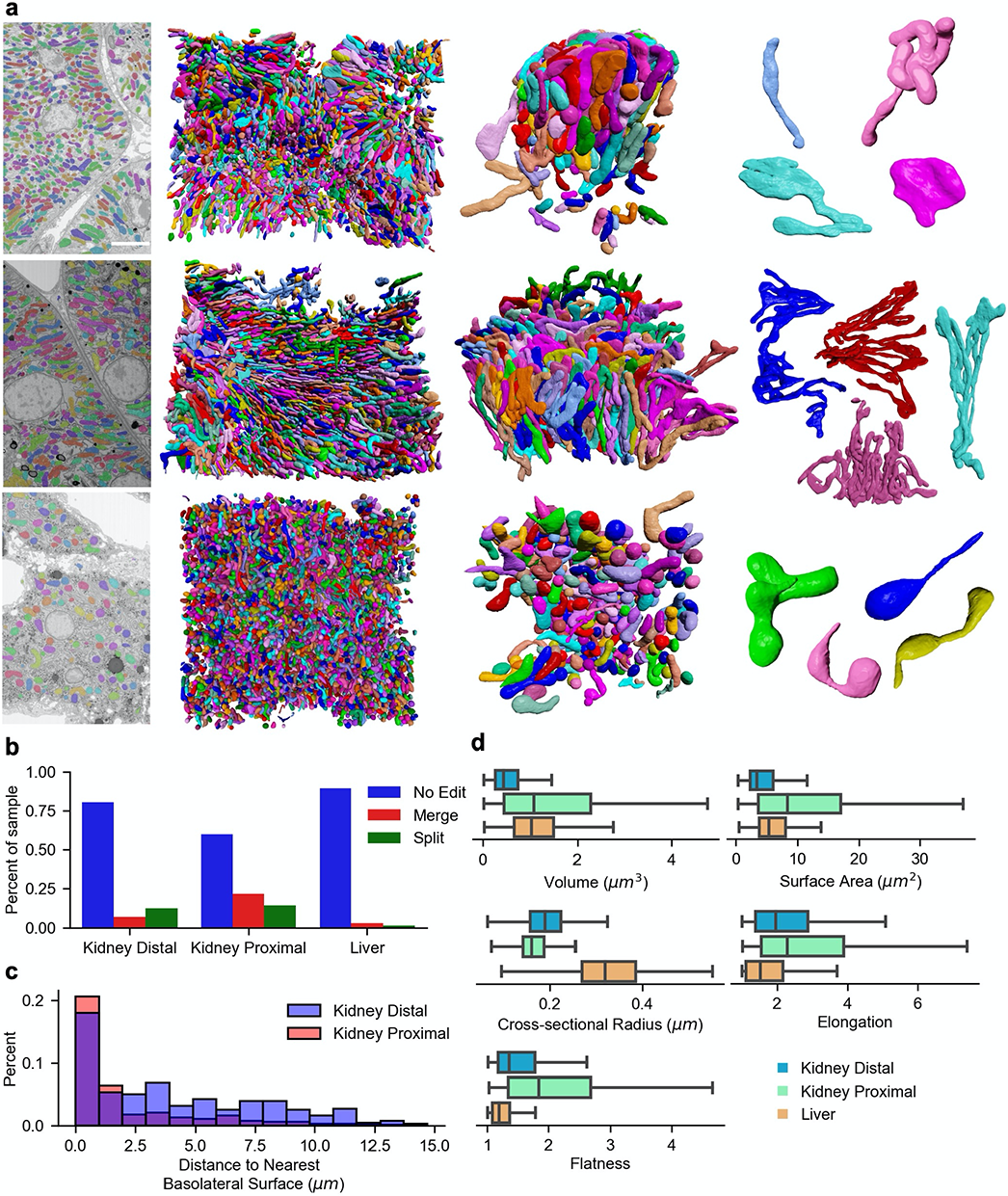
MitoNet results on volumes of mouse liver and kidney. **a.** Rows from top to bottom correspond to kidney distal tubule, kidney proximal tubule, and liver. Columns from left to right show: representative 2D images of MitoNet segmentation (scale bar, 5 μm), 3D predictions on the entire volume (small and boundary objects removed), a zoomed in ROI of raw model predictions (basolateral surface on top), representative mitochondrial models after manual cleanup. **b.** Plot of the fraction and type of cleanup operation required for a randomly chosen sample of model-predicted instances from kidney distal (n=347), kidney proximal (n=256) and liver (n=319) tissue. **c.** Plot of distance to the nearest basolateral surface, in microns, for randomly sampled mitochondria from kidney distal (blue) and kidney proximal (red) volumes after cleanup. **d.** Comparison of mitochondrial volume, surface area, cross-sectional radius, elongation, and flatness across the three volumes after cleanup (outliers not shown). Blue, kidney distal (n=405), green, kidney proximal (n=250), orange, liver (n=321).

Notably, 89%, 80%, and 60% of randomly sampled instances from the hepatocyte, distal tubule, and proximal tubule volumes, respectively, required no edits. In line with our results on the benchmarks, MitoNet predictions rarely required per-pixel corrections, but merge and split errors occurred in a data-specific manner (Figure 5b). For example, 12% of sampled distal tubule instances required splitting, while 21% in the proximal tubules needed merging – multiple merge operations were often required for complex mitochondria. With empanada’s point-and-click proofreading operations we took under 2 hours to split and merge all of the sampled distal tubule instances. Proofread instance segmentations can be used for a variety of quantitative analyses. We assessed mitochondrial polarization by measuring distance from basolateral surfaces in distal and proximal tubules (Figure 5c) and discovered statistically significant differences (median distance in distal tubule, 3.38 μm; proximal tubule, 0.86 μm, p<0.001). Additionally, other simple measurements accessible only with instance segmentation, e.g., volume, surface area, cross-sectional radius, elongation, and flatness, also showed intriguing differences between these groups (Figure 5d, statistical measures summarized in Table 3). Such large-scale observations of mitochondrial architectures in 3D and at high resolutions may reveal new and unexpected insights in kidney and other tissue pathologies.

**Table 3.** P-values for mitochondrial parameters measured between samples from mouse kidney and liver datasets. Diagonal entries compare the p-values calculated for a cleaned up random sample against unmodified model output. Bolded entries are statistically significant at a threshold of 0.05.

|  | Volume<br>(Mann-Whitney U) |  |  | Surface Area<br>(Mann-Whitney U) |  |  | Cross-sectional Radius<br>(Student's T-test) |  |  | Elongation<br>(Student's T-test) |  |  | Flatness<br>(Student's T-test) |  |  |
| --- | --- | --- | --- | --- | --- | --- | --- | --- | --- | --- | --- | --- | --- | --- | --- |
| Dataset | Kidney<br>Distal | Kidney<br>Proximal | Liver | Kidney<br>Distal | Kidney<br>Proximal | Liver | Kidney<br>Distal | Kidney<br>Proximal | Liver | Kidney<br>Distal | Kidney<br>Proximal | Liver | Kidney<br>Distal | Kidney<br>Proximal | Liver |
| Kidney<br>Distal | **<0.001** | **<0.001** | **<0.001** | **<0.001** | **<0.001** | **<0.001** | **0.018** | **<0.001** | **<0.001** | 0.906 | **<0.001** | **<0.001** | **0.006** | **<0.001** | **<0.001** |
| Kidney<br>Proximal |  | 0.141 | 0.244 |  | 0.349 | **<0.001** |  | **0.001** | **<0.001** | **<0.001** | 0.764 | **<0.001** |  | 0.146 | **<0.001** |
| Liver |  |  | **0.008** |  |  | **0.027** |  |  | 0.151 | **<0.001** | **<0.001** | 0.893 |  |  | 0.523 |

## Discussion

In this paper we have reported several resources for the growing volume EM community: a massive and heterogeneous unlabeled cellular EM image dataset that samples over 2 PB of EM data (CEM1.5M), a similarly heterogeneous dataset of >135,000 instances of labeled mitochondria (CEM-MitoLab), a model trained on this dataset (MitoNet), a set of diverse benchmarks (MitoNet Benchmarks) for model testing, and a Python package and napari plugin (empanada) for immediate use of MitoNet with proofreading tools. We also describe these datasets with a simple spreadsheet implementation of REMBI ^55^ containing image and biological metadata. Together, these resources relieve the “segmentation bottleneck”, a current constraint in volume EM workflows otherwise replete with technological and application advances. Crucially, CEM1.5M and empanada can be used for other features in cellular EM, and while CEM-MitoLab, MitoNet and benchmarks are mitochondria specific resources, their strong performance validates the strategy of broad but shallow sampling of cellular contexts to create general segmentation models for any organelle. Heterogeneity of training data is crucial, as shown by the surprising result that models trained on our noisy and inaccurate crowdsourced annotations generalize better than models trained on narrow but expertly labeled data like MitoEM. Expert proofreading was still necessary to achieve our best results, so broader expert participation in future shared efforts could greatly benefit the field. Similarly, homogeneous volume EM benchmarks derived from connectomics data are poor tests of generalization (indeed, the Lucchi++ benchmark has been mined to the point of performance saturation ^49, 56^). The assortment of mitochondria, contexts, quality, and overall difficulty in our benchmarks offer a stiff test for new models, but future benchmarks must be continually expanded to avoid exhaustion.

We made an important decision to forego 3D approaches. This allowed us to leverage abundant 2D datasets to represent a much broader range of cells and tissues, and also maximized images sampled – our crowdsourced dataset of ∼6,000 2D labeled images would have equated to just 30 3D labeled images of comparable dimensions. Staying in the 2D regime also relieves computational burden (although our workflows can work both on consumer-grade laptops without GPUs, or on HPC clusters with many), congruent with our belief that true democratization demands that such tools be usable without DL expertise or expensive compute resources. The methods we developed in this work, such as median filtering and ortho-plane inference, retain the efficiencies of labeling and running model inference in 2D but utilize 3D context where possible. Eventually, we expect native 3D approaches to surpass our methods. Even then, empanada can be used to quickly segment and proofread the 3D data needed to train this next generation of models. For mitochondrial models in 2D or 3D, MitoNet can provide high-quality initial predictions.

True human-level mitochondrial instance segmentation on any 2D or 3D EM image may be possible. Better methods are needed to segment small and closely packed instances, and low contrast and resolution or unusual cellular contexts currently pose difficulties to MitoNet. That said, semantic segmentation quality was extremely strong on most datasets tested, and a proof-of-principle experiment on kidney and liver volumes shows that MitoNet is already a powerful and accessible tool for rapid visualization and accurate quantification of mitochondrial morphologies, even without proofreading for some datasets (Table 3). Current DL pipelines rely on repeated cycles of human-in-the-loop annotate, train, infer, and proofread, as required by specialist models. Generalist DL segmentation models, like MitoNet for mitochondria, reduce the cycle to infer directly and proofread on any volume EM dataset. Even for datasets where out-of-the-box inference is poor, annotation by sparsely labeling 2D patches is quick, and allows finetuning with trivial compute time. Future expansions of the CEM resources will further enable model generality irrespective of feature segmented, and such models, whether for semantic, instance, or panoptic segmentation, can be trained using empanada and broadly shared for deployment in the napari plugin.

Volume EM is catalyzing a “quiet revolution” ^57^ by enabling large-volume high-resolution 3D images of cellular and subcellular features, revealing insights into connectomics and cell biology. Accurate and efficient extraction of features of interest from these massive ultrastructural images lends itself to computational solutions. Here we create large-scale and relevant training data resources and train a generalist model to segment mitochondria in any EM image. In tandem with recent initiatives to generate and share large volume EM reconstructions ^36^, our methods provide a blueprint for future applications, thereby expanding the segmentation toolkit and helping accelerate discoveries in this exciting field.

## Materials and Methods

### CEM1.5M Creation

The expansion of CEM500K to create the CEM1.5M dataset followed the data standardization and curation protocols presented in our previous work ^35^. External datasets were either downloaded in their entirety or, for large datasets stored online in next generation file formats (n5 or zarr), accessed either with the CloudVolume or fibsem-tools APIs. Datasets larger than 5 GB were randomly cropped into 320 cubes of 256^3^. Metadata and proper attribution for each dataset is available in **Supplementary file 1.** TEM and STEM images were grouped into directories based on imaging project or publication before processing. 3,000 NCI TEM images with magnifications between 1000x (∼16 nm pixel) and 3000x (∼6 nm pixel) and excluding negative stain and immunogold label images were randomly selected from a library of over 2×10^5^ images. Metadata for these images was unavailable.

### Crowdsourced Annotation

Our Zooniverse workflow closely approximated the Etch-a-cell project ^13^. Flipbooks of image patches from 3D data (8-bit tif stacks of five consecutive images of 224×224) were contrast rescaled to 25-235 and interpolated to 480×480 for easier viewing. For 2D images contrast was similarly rescale but crops of 512×512 were used. Over the course of the project, the Zooniverse chat channel and a formal instruction session were used to review examples and explain errors. During expert proofreading, the correction of FP and FN detections was emphasized over pixel-level painting; disagreements were discussed and resolved as a group. All annotations were passed through a connected components filter after proofreading.

The input to the consensus algorithm was a set of **N** annotations (the retirement limit) containing **K** mitochondrial detections (Supplementary Figure 3 i). An undirected graph was initialized where each node corresponded to one of the **K** detections. Edges were added to the graph to connect detections with mask IoU scores that exceeded a small threshold value of 0.1 (the same algorithm, when applied during ortho-plane inference, used a threshold of 0.01). Each connected component in the detection graph was then processed independently. Nodes in a connected component were organized into cliques where all detections within a clique shared edges with IoU scores greater than 0.75 (Supplementary Figure 3 ii,iii). Edges between detections in different cliques were retained. An iterative algorithm was applied to determine whether cliques should be merged or remain split. First, the most connected clique (i.e., the one with the most edges; node C in Supplementary Figure 3 ii,iv; A in 3 iii,v) was selected. Second, if any of the most connected clique’s neighbors contained more detections, then the most connected clique was dissolved and its detections were pushed out to each of its neighbors (C in Supplementary Figure 3 iv). Otherwise, all the most connected clique’s neighbors were dissolved and their detections were merged into the most connected clique (B and D in Supplementary Figure 3 iv, B and C into A in 3 v). These two steps were repeated until no edges were left between cliques. Each clique represented an object instance. The detection masks within a clique were added together to form an image where each pixel had a value from 1 to **N**. A vote threshold was applied to create the final binary instance mask. All instances were combined into the final consensus annotation (Supplementary Figure 3 vi).

### Labeled and benchmark dataset creation

The “legacy” portion of CEM-MitoLab came from publicly available mitochondrial segmentations (Kasthuri++, Guay, UroCell, MitoEM, Heinrich et al. and Perez et al.) ^37, 58, 59^ and previous in-house segmentation projects. Of these, MitoEM already had instance labels, and a connected components filter was applied to convert other semantic labels to instance segmentations. All in-house segmentation projects were proofread or annotated by experienced researchers. The projects included annotations of whole volumes, TEM images, and a random selection of patches from CEM500K ^35^; none were derived from the benchmark datasets. For volumes and TEM images, 512×512 crops were taken from xy planes, or all three planes for isotropic volumes, and passed through the deduplication and filtering pipeline to remove redundant and uninformative patches. Legacy images were used as-is except for MitoEM-H and MitoEM-R volumes which were binned to 16 nm pixels and Heinrich et al. images which were downloaded at 8 nm resolution. To create the alternative training datasets in Figure 4, MitoEM and Heinrich et al. ground truth ROIs were sliced to 2D images. For MitoEM, patches of size 512×512 were cropped from xy planes only. The Heinrich et al. volumes were cropped into patches of 224×224 or smaller from xy, xz and yz planes, and 2D segmentations passed through a connected components filter.

The HeLa cell, C. elegans, fly brain and salivary gland benchmark datasets were annotated by ariadne (https://ariadne.ai/). The glycolytic muscle ^32^ dataset was annotated in-house. The benchmark derived from Lucchi++ was generated by applying a connected components filter to the binary mitochondrial labelmaps followed by manual proofreading. To be rigorous, we excluded from CEM1.5M and CEM-MitoLab not just the benchmark datasets, but also images related to them, e.g., the entire Lucchi++ training dataset, all other drosophila neuronal data from OpenOrganelle, and FIB-SEM and STEM data from the in-house C. elegans project.

### CEM1.5M Pre-training

Unsupervised pre-training used the SwAV algorithm ^48^. A ResNet50 ^47^ model was trained for 200 epochs with a batch size of 256; other hyperparameters used defaults defined in https://github.com/facebookresearch/swav. Image augmentations included 360-degree rotations, randomly resized crops, brightness and contrast jitter, random Gaussian blur and noise, and horizontal and vertical flips. To correct for the imbalance in the number of patches per dataset, weighted random sampling was applied. Weights per dataset were calculated by:

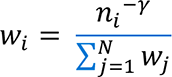

*γ* was a float from 0 to 1, *n_i_* was the number of patches from the *i^th^* dataset, and N was the total number of datasets. We used *γ*=0.5.

### Model architecture, training parameters, and post-processing

MitoNet was based on Panoptic-DeepLab (PDL). PDL is an encoder-decoder style network that uses atrous spatial pyramid pooling (ASPP) to integrate features over multiple scales; our configuration included one ASPP decoder for semantic segmentation prediction and another for instance center and offsets prediction. A ResNet50, pre-trained as detailed above, was used as the encoder network. To preserve spatial information, strides in the last downsampling layer were replaced with dilation such that the output from the encoder was 16x smaller than the input. Dilation rates in the ASPP modules were set to 2, 4, and 6 with 256 channels per convolution and dropout probability of 0.5. The semantic and instance decoders had a single level at which the output from the first layer in the encoder was fused to the interpolated ASPP output via depthwise separable convolution. Convolutions had 32 channels in the semantic decoder and 16 in the instance decoder. The semantic segmentation, instance center and offset heads used a single depthwise separable convolution with kernel size of 5. The PointRend ^44^ module was added to refine the semantic segmentation. Module parameters used the defaults defined in Detectron2 ^60^. Briefly, during training, the segmentation logits were upsampled by a factor of 4 to the original image resolution and then refined in a single step. During evaluation, two refinement steps were applied with the segmentation logits being upsampled by a factor of 2 at each step. An arbitrary number of interpolation and refinement steps can be applied sequentially to upsample the segmentation to a desired resolution. The overall loss function was:

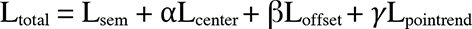

α, β, *γ* were constants set to 200, 0.01 and 1, respectively. L_sem_ was the semantic segmentation loss computed as the bootstrapped (binary) cross-entropy where only the top 20% of cross-entropy values were averaged across a batch. L_center_ was the instance center regression loss and L_offset_ was the center offset loss calculated as mean squared error and absolute error (L1), respectively. L_pointrend_ was the PointRend loss calculated as the (binary) cross-entropy. MitoNet was trained on CEM-MitoLab using the One Cycle learning rate policy ^61^ with AdamW ^62^ for 120 epochs. The max learning rate was set of 0.003 with weight decay of 0.1 and momentum cycled from 0.85 to 0.95. Learning rate warmup lasted for the first 30% of training epochs. Image augmentations included large scale jitter ^63^, random cropping of a 256×256 patch, 360-degree rotations, brightness and contrast adjustments, and vertical and horizontal flips. Weighted sampling – as defined above for CEM1.5M – used *γ*=0.3.

Postprocessing of the model outputs in an instance segmentation used the method developed for Panoptic-DeepLab. In all experiments, the non-maximum suppression kernel size was set to 7 and the center confidence threshold was set to 0.1.

### Benchmark inference and evaluation

Ortho-plane inference ^46^ was used on all benchmark volumes. During inference over each plane, median semantic segmentation probabilities were calculated from a queue of a few consecutive slices. A queue length of 3 was used for the C. elegans and fly brain, 5 for the HeLa and glycolytic muscle and 7 for the Lucchi++ and salivary gland. Queue lengths roughly track inversely with voxel size. Median probabilities were hardened at a confidence threshold of 0.3 and Panoptic-DeepLab postprocessing was performed. The resultant instance segmentations were passed through a connected components filter. After forward and backward instance matching with IoU and IoA threshold of 0.25, a size and bounding box extent filter were applied to eliminate likely false positives. Minimum sizes were 500 voxels for C. Elegans and fly brain; 800 for HeLa; 1,000 for salivary gland; 3,000 for glycolytic muscle. Minimum bounding box extent was fixed at eight voxels. The same consensus segmentation algorithm as above was applied to ensemble the three segmentation stacks created by ortho-plane inference. A clique IoU threshold of 0.75 and a vote threshold of 2 out of 3 was used for all benchmarks except the C. elegans and salivary gland volumes. For those a vote threshold of 1 out of 3 was used. This was in response to the relatively low IoU scores observed. Importantly, while a vote threshold of 1 was used, each clique was required to have at least 2 detections; this enabled increases in IoU score without the introduction of many FPs. The consensus algorithm occasionally produced overlapping instances. Instances overlapping more than 100 voxels were merged; otherwise, the few overlapping voxels would be assigned to whichever instance was processed last.

### OpenOrganelle volume inference and analysis

For the “Mouse kidney” dataset, two ROIs containing cells in the proximal tubule and distal tubule exclusively (some endothelial cells could not be avoided) were manually chosen from the 128nm overview to download the desired data at 16nm resolution. One ROI from the “Mouse liver” dataset was similarly selected. Inference used the best version of the MitoNet model and was run only on slices from the xy plane with a segmentation confidence threshold of 0.3 and median filter size of 5. A single compute node with a P100 GPU and 16 GB of memory was used. Image volumes and predicted mitochondrial instances were downsampled to 32nm pixel size for visualization and cleanup and a size threshold of 1,000 voxels was applied. For cleanup and quantification, 400 instances were randomly chosen such that they did not touch the boundary and each bounding box was at least 15 pixels long on a side. The empanada napari plugin was used to proofread each instance and merge, split, paint, and erase as required. FPs, mitochondria in endothelial cells, and instances on the boundary after merging were removed. Volume, surface area, flatness and elongation measurements for individual mitochondria were calculated using LabelShapeStatistics filter as implemented in ITK ^64^ and cross-sectional radius was calculated as described in **Supplementary Methods**. Basolateral surfaces were approximated by fitting a triangular mesh to manually placed periodic fiducials using the ball pivoting algorithm ^65^ and the minimum distances between vertices of each mitochondrial mesh to the nearest basolateral surface were calculated.

## Supporting information

Supplementary File 2: MitoNet performance against benchmarks

Supplementary File 1: metadata for CEM-MitoLab

Supplementary Movie: Instance segmentation by MitoNet on large kidney and liver volumes

## Acknowledgements

This project has been funded in whole or in part with Federal funds from the National Cancer Institute, National Institutes of Health, under Contract No. 75N91019D00024. The content of this publication does not necessarily reflect the views or policies of the Department of Health and Human Services, nor does mention of trade names, commercial products, or organizations imply endorsement by the U.S. Government. This publication uses data generated via the Zooniverse.org platform, development of which is funded by generous support, including a Global Impact Award from Google, and by a grant from the Alfred P. Sloan Foundation. We thank Helen Spiers, Martin Jones and Lucy Collinson for help with Zooniverse. We thank the WHK-SIP program and batch of 2021 student annotators, especially student leaders Ella Fitzgerald and Taeeun Kim. Valentina Baena, Adam Harned, Kunio Nagashima and Heather Berensmann helped perform proofreading. Patrick Friday helped curate metadata, and Aayush Bhatawadekar contributed to renderings and plugin development. Finally, we appreciate the volume EM community, especially the Data Working Group for general discussions. This work utilized the computational resources of the NIH HPC Biowulf cluster. (http://hpc.nih.gov).

## Data and Code Availability

All resources are linked with additional explanation on our project webpage https://volume-em.github.io/empanada. Code for the empanada library is available at https://github.com/volume-em/empanada. Code for the empanada-napari plugin is available at https://github.com/volume-em/empanada-napari. All datasets are accessible through EMPIAR: CEM1.5M (https://www.ebi.ac.uk/empiar/EMPIAR-11035/), CEM-MitoLab (https://www.ebi.ac.uk/empiar/EMPIAR-11037/), MitoNet Benchmarks (https://www.ebi.ac.uk/empiar/EMPIAR-10982/). Pre-trained models are available on Zenodo: CEM1.5M weights (https://zenodo.org/record/6453160#.YnPjEy-cbTR), MitoNet (https://zenodo.org/record/6327742#.YnLciy-cbTS).

---

**Supplementary Figure 1:**
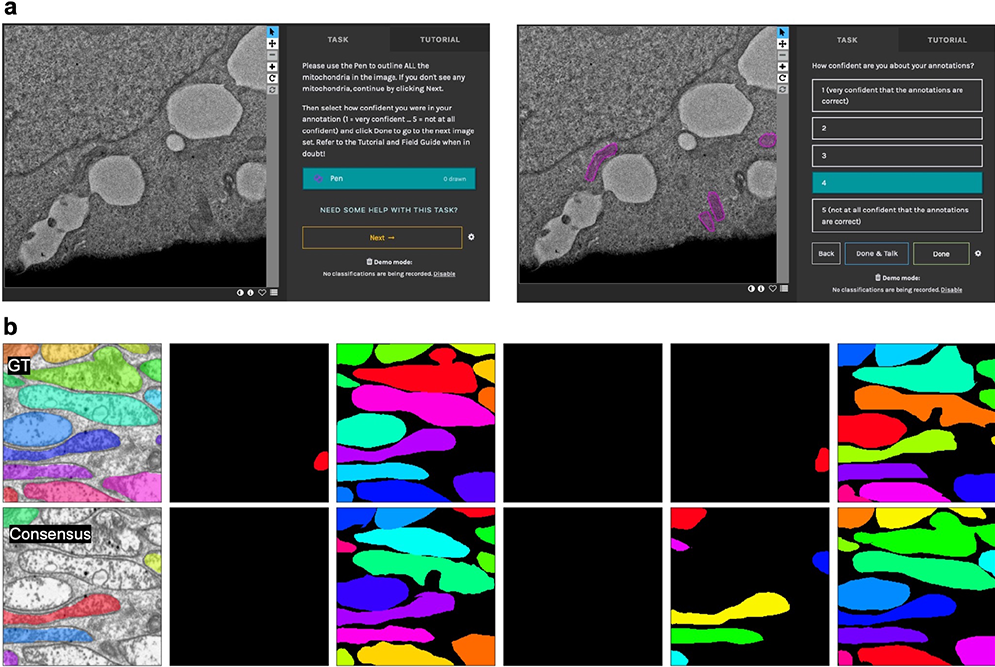
**a.** Screenshots from the Zooniverse project showing the user annotation interface (left) and example annotation with confidence rating (right). **b.** Example of crowdsourced annotation with ground truth (top left), consensus annotation (bottom left) and ten independent student annotations of an image showing a low degree of consensus; the all-or-nothing pattern suggest differences in annotator knowledge or experience.

**Supplementary Figure 2:**
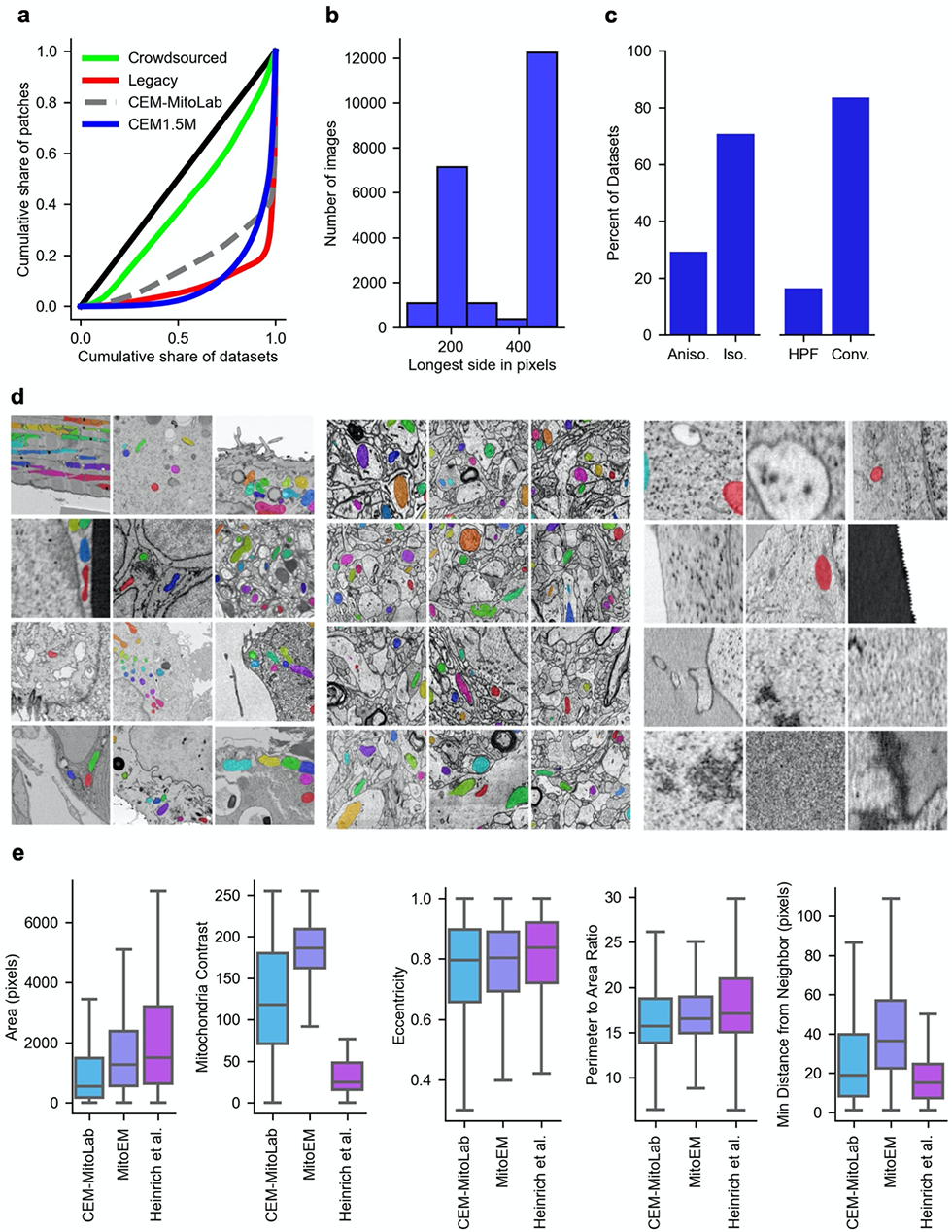
**a.** Lorenz plots for CEM1.5M (blue, Gini coefficient = 0.802), crowdsourced data (green, 0.209), legacy data (red, 0.840), CEM-MitoLab (dashed gray, 0.686). Black line, perfect equality in distribution (Gini = 0). **b.** Distribution of longest image side in pixels for patches in CEM-MitoLab. **c.** Percentage of isotropic versus anisotropic 3D data (left, n=489) and sample fixation method (right, n=593) in source datasets that make up CEM-MitoLab. **d.** Random selection of image patches from CEM-MitoLab (left), Mito-EM (middle) and Heinrich et al. (right). **e.** Comparison between 2D mitochondrial instance label maps in CEM-MitoLab (blue), MitoEM (purple) and Heinrich et al. (magenta) labeled datasets, with respect to (left to right) area in pixels, contrast, eccentricity, perimeter to area ratio, and distance from nearest neighbor in pixels.

**Supplementary Figure 3.**
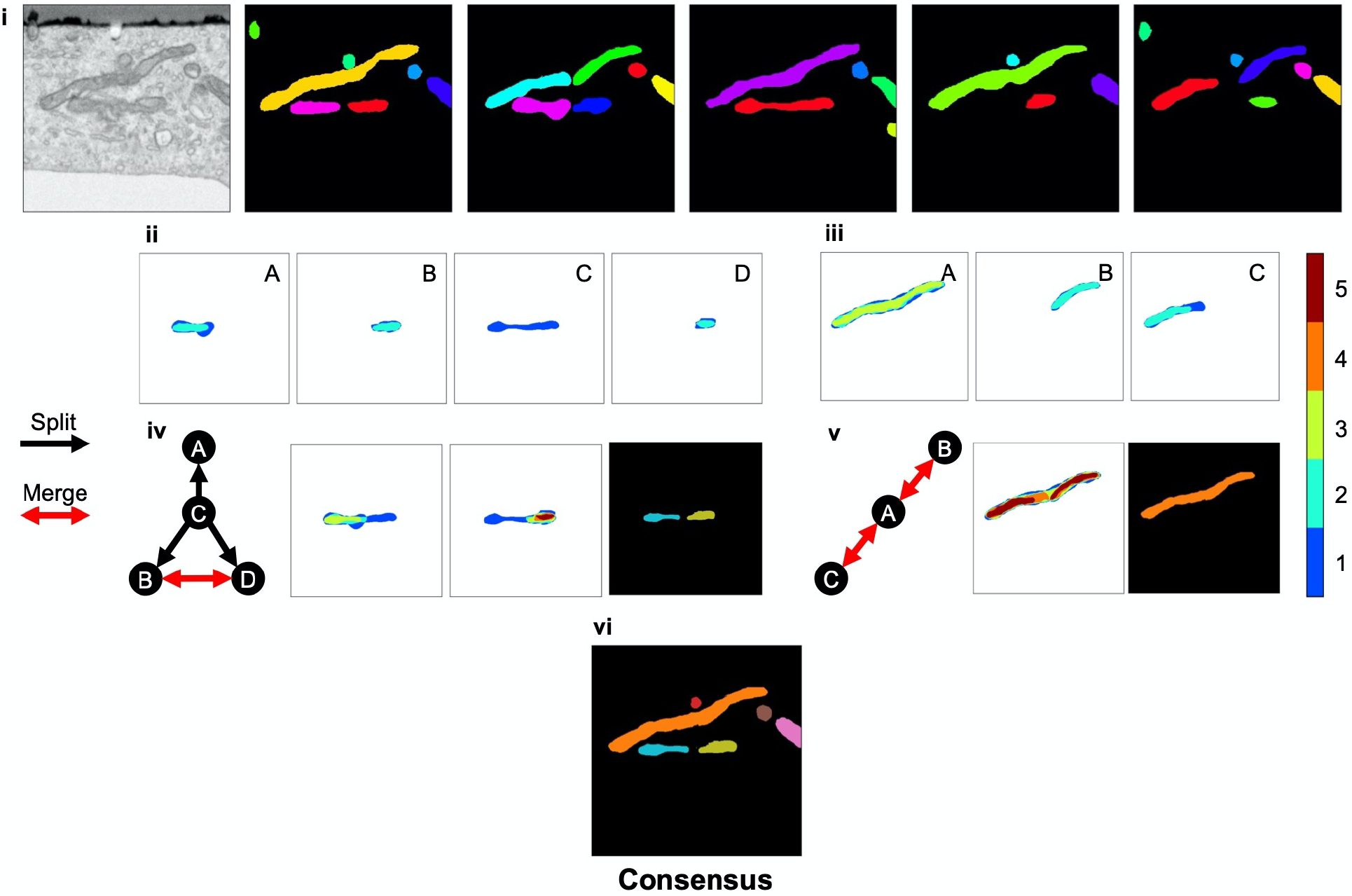
Example of consensus algorithm splitting and merging instance votes to create accurate consensus label maps. See Materials and Methods for more details. Colorbar (right) represents number of votes per pixel.

**Supplementary Figure 4.**
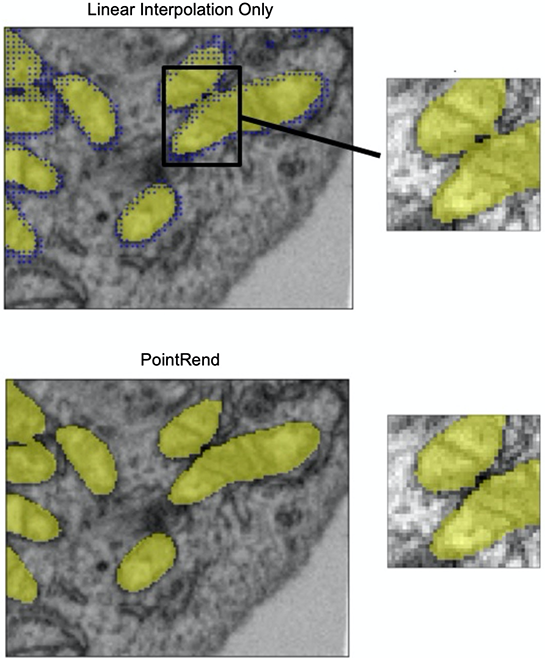
Example image showing difference in semantic segmentation borders of closely apposed mitochondria, after linear interpolation (top) or PointRend (bottom). Blue dots, PointRend sampling locations.

**Supplementary Figure 5.**
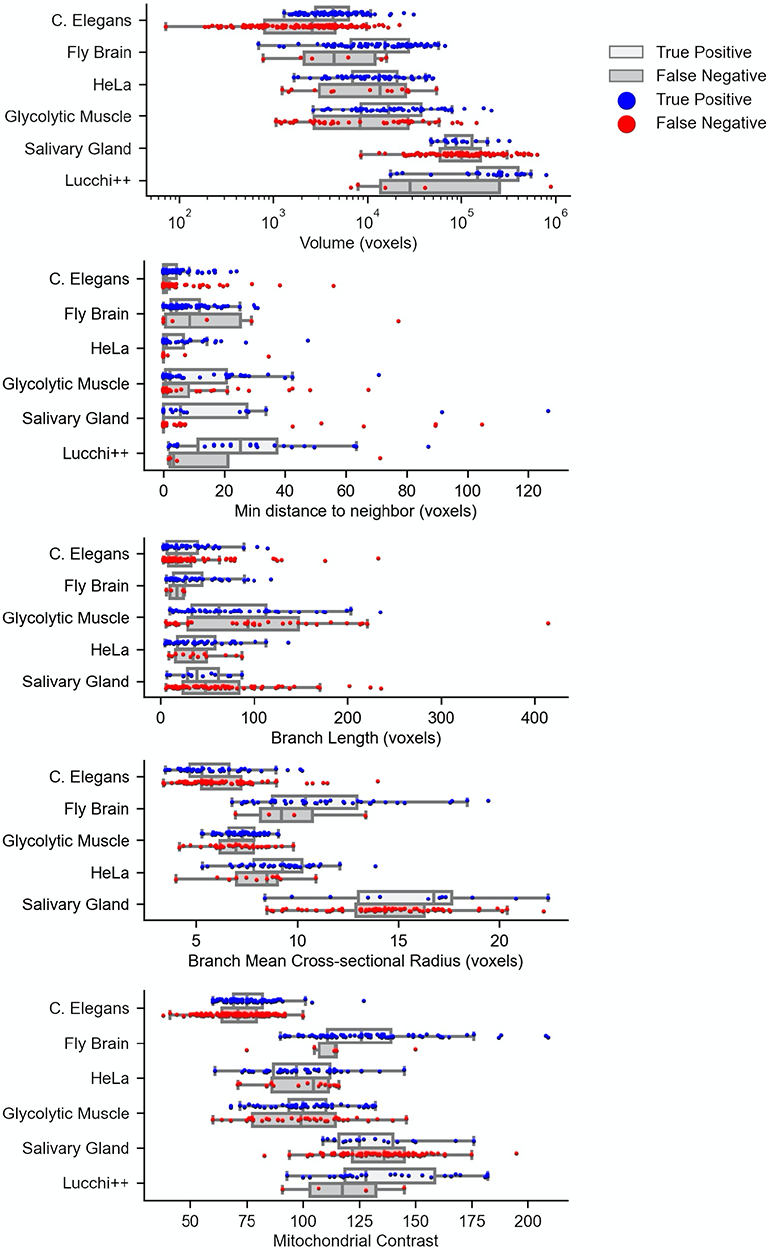
Analysis of True Positive (TP) and False Negative (FN) detections by MitoNet on benchmarks, grouped by different mitochondrial attributes. Top to bottom: mitochondrial volume (log scale), minimum distance to nearest neighbor, branch length, branch mean cross-sectional radius (measurements by voxel) and mitochondrial contrast. TP and FN detections were calculated at an IoU threshold of 0.5. Branch length and cross section plots exclude the Lucchi++ benchmark, as the branches for FN mitochondria were shorter than the skeleton pruning threshold. Finetuned model results are plotted for salivary gland.

**Supplementary Figure 6.**
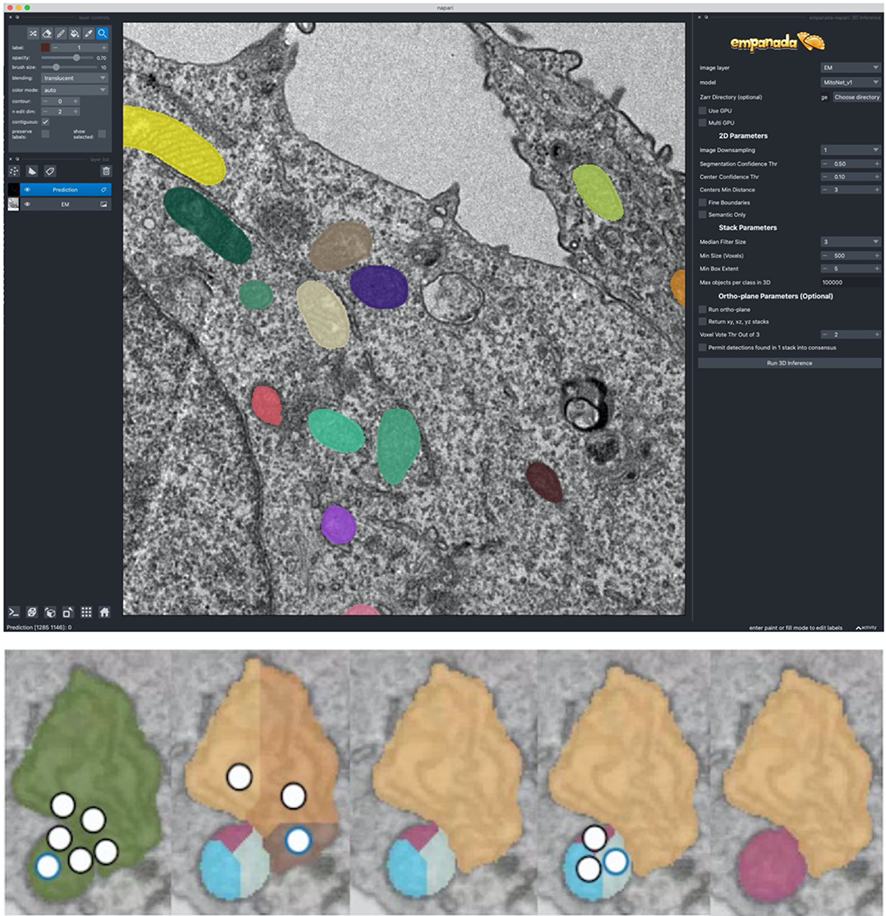
Top, screenshot of the empanada plugin in napari. Bottom, Illustration of split and merge operations by manual placement of fiducials in the plugin.

**Supplementary Table 1.** The effect of postprocessing steps on benchmark performance. The numbers in bold are the best IoU and F1@75 scores.

|  | 2D Stack Only |  | IoA Merging |  | Median Filtering |  | Ortho-plane<br>Vote Thr. of 1 |  | Ortho-plane<br>Vote Thr. of 2 |  |
| --- | --- | --- | --- | --- | --- | --- | --- | --- | --- | --- |
| Dataset | IoU | F1@75 | IoU | F1@75 | IoU | F1@75 | IoU | F1@75 | IoU | F1@75 |
| C. Elegans | 0.473 | 0.223 | 0.512 | 0.210 | 0.543 | 0.251 | **0.604** | **0.325** | 0.439 | 0.183 |
| Fly Brain | 0.876 | 0.754 | 0.877 | 0.790 | 0.879 | 0.821 | 0.892 | **0.890** | **0.902** | 0.855 |
| HeLa | 0.690 | 0.178 | 0.690 | 0.203 | 0.708 | 0.308 | 0.784 | **0.636** | **0.791** | 0.503 |
| Glycolytic Muscle | 0.796 | 0.225 | 0.829 | 0.508 | 0.835 | 0.519 | 0.805 | 0.450 | **0.839** | **0.601** |
| Salivary Gland | 0.041 | 0.000 | 0.048 | 0.000 | 0.056 | 0.000 | **0.103** | **0.008** | 0.035 | 0.000 |
| Lucchi++ | 0.851 | 0.659 | 0.852 | 0.727 | 0.859 | 0.817 | 0.862 | 0.800 | **0.871** | **0.879** |

**Supplementary Table 2.** The effect of model pre-training on benchmark performance. The self-supervised algorithm used was either momentum contrast (MoCov2) with the previous iteration of the CEM dataset, CEM500K, or SwAV, with either unbalanced (*γ*=0) or balanced (*γ*=0.5) sampling from CEM1.5M.

| | No Pre-training | | CEM500K-moco | | CEM1.5M-SwAV<br>Unbalanced ($\gamma=0$) | | CEM1.5M-SwAV<br>Balanced ($\gamma=0.5$) | |
| --- | --- | --- | --- | --- | --- | --- | --- | --- |
| Dataset | IoU | F1@75 | IoU | F1@75 | IoU | F1@75 | IoU | F1@75 |
| C. Elegans | **0.666** | 0.288 | 0.638 | **0.332** | 0.622 | 0.280 | 0.604 | 0.325 |
| Fly Brain | 0.899 | 0.896 | 0.897 | 0.886 | **0.910** | **0.916** | 0.902 | 0.885 |
| HeLa | 0.802 | 0.467 | **0.817** | 0.500 | 0.784 | 0.453 | 0.791 | **0.503** |
| Glycolytic Muscle | **0.848** | 0.558 | 0.839 | 0.561 | 0.832 | 0.532 | 0.840 | **0.601** |
| Salivary Gland | **0.315** | **0.013** | 0.171 | 0.000 | 0.130 | 0.000 | 0.103 | 0.008 |
| Lucchi++ | **0.880** | 0.844 | 0.882 | 0.844 | 0.864 | 0.813 | 0.871 | **0.879** |
| Average | **0.735** | 0.511 | 0.707 | 0.521 | 0.690 | 0.499 | 0.685 | **0.534** |

**Supplementary Table 3:** Average mitochondrial measurements split by true positive (TP), false negative (FN) with corresponding p-values across benchmarks. Bolded entries are statistically significant at a threshold of 0.05.

|  | Volume (voxels) |  |  | Mitochondrial Contrast |  |  | Min. distance to neighbor (voxels) |  |  | Branch length (voxels) |  |  | Branch mean cross-sectional radius (voxels) |  |  |
| --- | --- | --- | --- | --- | --- | --- | --- | --- | --- | --- | --- | --- | --- | --- | --- |
| Dataset | TP | FN | P | TP | FN | P | TP | FN | P | TP | FN | P | TP | FN | P |
| C. Elegans | 5489.9 | 3504.0 | **<0.001** | 75.9 | 70.6 | **<0.001** | 3.2 | 2.2 | 0.253 | 26.6 | 31.2 | 0.481 | 5.8 | 6.3 | 0.150 |
| Fly brain | 19177.5 | 6944.0 | 0.051 | 128.6 | 112.3 | 0.132 | 7.8 | 20.6 | **0.004** | 34.9 | 16.7 | 0.201 | 11.1 | 0.7 | 0.389 |
| HeLa | 15271.2 | 19072.2 | 0.330 | 100.3 | 95.9 | 0.361 | 5.2 | 2.2 | 0.238 | 41.7 | 43.0 | 0.891 | 9.1 | 7.9 | **0.030** |
| Glycolytic Muscle | 34437.6 | 19837.0 | **0.033** | 100.6 | 99.0 | 0.676 | 12.6 | 6.4 | **0.039** | 78.9 | 94.7 | 0.298 | 7.3 | 7.0 | 0.223 |
| Salivary Gland | 113909.7 | 133460.1 | 0.462 | 130.9 | 133.7 | 0.533 | 20.5 | 5.3 | **0.008** | 44.3 | 62.9 | 0.222 | 15.6 | 14.4 | 0.203 |
| Lucchi++ | 295145.0 | 177141.3 | 0.207 | 139.4 | 117.7 | 0.051 | 30.3 | 13.1 | 0.074 | N/A | N/A | N/A | N/A | N/A | N/A |

**Supplementary Movie 1. Mitochondrial instance predictions by MitoNet, in liver, kidney distal tubule and kidney proximal tubule volumes.** 2D predictions in volume EM stacks and 3D reconstructions are shown (with small objects and objects on boundaries removed), as well as examples of highly complex mitochondrial instances.

**Supplementary File 1. Spreadsheet of available metadata of in-house and externally sourced datasets contributing to CEM1.5M.**

**Supplementary File 2. Model performance scores, when trained by various labeled datasets, for individual benchmarks. The average of these benchmark scores is shown in** Fig 4d**. Salivary Gland benchmark required finetuning and was not included.**

