## Supplementary material for "Instance segmentation of mitochondria in electron microscopy images with a generalist deep learning model": Supplemetary Methods

*Zooniverse workflow*

Our Zooniverse workflow was greatly aided by the creators of, and thus closely approximated, the Etch-a-cell project^1^. Patches in the CEM dataset extracted from vEM images were reconstructed into short “flipbooks” (tif stacks of five consecutive images of 224x224) where each patch from the CEM dataset was the center image (the adjacent patches were typically filtered out by the CEM deduplication pipeline). Before uploading to Zooniverse each flipbook had its 8-bit intensity range rescaled from 25 to 235 and was interpolated to a size of 480x480 for easier viewing. For 2D images, crops of 512x512 were used. Intensity was rescaled to the same range as the flipbooks but resizing was unnecessary.

The first two sets of data uploaded to Zooniverse (Group 1a) had an annotation retirement limit of five (i.e., every image was annotated independently by five people) and were annotated by a mixture of lab members familiar with cellular EM and inexperienced student annotators. Consensus annotations from these sets only used annotations from experts; the students’ feedback on the project helped to refine the Zooniverse tutorials and interface. For all remaining sets, which were annotated entirely by an expanded cohort of students, the retirement limit was set to 10. For the first two of these sets, 5-10% of images were annotated by a team of 2-3 EM experts. These “gold-standard” annotations were used to measure the performance of the student annotators while the set of images was still actively being worked on and to provide the students with training examples from which to learn. A Google CoLab notebook was used to share the gold standard annotations with students. Later, as the students’ annotations became more accurate, only the consensus annotations were shared. In addition to this feedback, a single instruction session was organized to review challenging examples and explain common mistakes after the conclusion of the first set of images.

*Consensus algorithm explanation*

This algorithm assumed that the most connected clique at each step roughly corresponded to a maximally merged instance. When one of this clique’s neighbors contained more detections, this implied that more votes were in favor of splitting than maximal merging. Pushing the most connected clique to its neighbors ensured that no detections were deleted before being counted towards the final pixel-level majority vote. Conversely, when the most connected clique had the most detections, this implied that more votes were in favor of maximal merging and therefore all detections in neighboring cliques were pulled into the most connected clique.

*Proofreading*

Once all images in a set were retired, a consensus strength and instance segmentation were calculated for each. The consensus strength was the average F1@50 score of each individual annotation relative to the consensus instance segmentation. Lower consensus strengths indicated greater disagreement between annotators. Next, all images and instance segmentations were ordered by consensus strength and stacked into a single tif file for later proofreading. If the set used flipbooks, then the images and their consensus instance segmentations were resized from 480x480 back to their original sizes. Padding was added as needed to ensure that all images in the stack had the same dimensions.

Typically, the single proofreading stack was split into chunks of 40-50 images/flipbooks such that multiple experts could perform proofreading in parallel. Proofreaders were instructed to emphasize the correction of false positive and false negative detections over fine-grained instance boundary corrections. Annotations in each chunk of images were verified by a second proofreader and disagreements were discussed and resolved by 2-3 proofreaders. 3D Slicer^2^ was used to visualize and correct annotations. The chunks of proofread images and annotations were again combined into a single large stack. Every image, or every third image for flipbooks, and corresponding instance segmentation was stored as a tiff image with any padding cropped out.

*Mitochondrial measurements*

For benchmarks and mouse liver and kidney datasets, mitochondrial volumes were calculated by trivially counting the number of voxels per instance. Minimum distances between neighboring mitochondria were calculated by computing distance transforms for each instance’s complement (i.e., set of all voxels not inside the instance) and storing the minimum value calculated at each voxel. Branch lengths and cross-sectional diameters were calculated by first skeletonizing^3^ each mitochondrion and computing its distance transform. Lengths were simply the number of voxels in each branch, and the mean cross-sectional diameter was the average of distance transform values that overlapped with the skeleton. Branches shorter than 60 nm were pruned. Mitochondria contrast was calculated as the difference between maximum and minimum intensity voxels in each instance.

*IoU and IoA instance matching*

Intersection-over-union (IoU) scores were calculated between instance bounding boxes on consecutive 2D slices. For each pair of instances with non-zero bounding box IoU, a mask IoU and IoA score was calculated. The Hungarian algorithm^4^ was then applied to the mask IoU scores to assign instances such that total IoU over all assignments was maximized. Unassigned instances, or assigned instances with IoU score less than a threshold value, were considered unmatched. Unmatched instances with IoA score greater than a threshold value were matched to the instance with which they shared the highest IoA. The remaining unmatched instances were assigned new labels in the stack. After forward matching for the entire stack, a backward pass of matching was run. No new labels were assigned in the stack during the reverse pass.
